# Sea stickleback genome reveals repeated chromosomal rearrangements in sticklebacks

**DOI:** 10.64898/2026.08.30.747834

**Authors:** Jule Drewalowski, Sergei Kliver, Leon Hilgers, Peter Rask Møller, Sarah S.T. Mak, Iva Kovačić, Bent Petersen, Joseph Nesme, Ann M. Mc Cartney, Alice Mouton, Giulio Formenti, Hannes Svardal, Genevieve Diedericks, Henrique G. Leitão, Rosa Fernández, Nuria Escudero, Judit Salces-Ortiz, Claudio Ciofi, Chiara Natali, Maria Angela Diroma, Alessio Iannucci, Marco Sollitto, Michael Hiller, M Thomas P Gilbert, Josefin Stiller

**Author notes:** Corresponding author: Josefin Stiller,; Jule Drewalowski.

## Abstract

Sticklebacks (Gasterosteidae) encompass model organisms which are of particular interest for evolutionary and ecological genomics. Within Gasterosteidae, chromosome number is variable (2n=40-46) and independent fusions of homologous chromosomes have been proposed. The sea stickleback (or fifteen-spined stickleback, *Spinachia spinachia*) has the lowest known number of chromosomes (2n=40) and hence is crucial in understanding chromosome evolution among sticklebacks, but is so far missing in genomic datasets. Here, we present a high-quality diploid genome assembly of *S. spinachia*. PacBio HiFi and Hi-C reads were assembled into a genome of 407.5 Mb in size, consisting of 20 chromosomes, with an N50 of 6.6 Mb and 98.96% complete single-copy BUSCO genes. A phylogenetic tree inferred across five stickleback species and four outgroup genomes from 19,156 genes, alongside synteny analyses and ancestral chromosome reconstructions, confirms *S. spinachia* as the sister species to the four-spined stickleback (*Apeltes quadracus*) and not as the sister group to all other sticklebacks as once thought. It has one species-specific chromosome fusion and shares two fusions with the three-spined stickleback (*Gasterosteus aculeatus*), none of which are present in its sister species. One of these fusions is also present in *Pungitius*, leading to reinterpretion of this fusion as ancestral to Gasterosteidae, with subsequent fission in *Apeltes*. This implies a lower ancestral chromosome number in Gasterosteidae (2n=44) than previously thought. The other fusion shared with *G. aculeatus* presents a case of convergence. Our results suggest that karyotype evolution in Gasterosteidae has been shaped by ancestral chromosome fusion, convergent fusion, and secondary fission.

## Introduction

Sticklebacks (family Gasterosteidae, Cottoidei) are one of the most important model organisms for studying adaptive evolution. In particular, the three-spined stickleback (*Gasterosteus aculeatus*) has emerged as a key model organism, with extensive genomic resources enabling studies of speciation and adaptation across diverse ecological contexts (McKinnon and Rundle 2002; Gibson 2005; Reid et al. 2021). These studies have provided important insights into processes such as repeated adaptation to freshwater environments and the genetic basis of associated phenotypic evolution, including distinct ecotypes and behavioral differences (Colosimo et al. 2005; Kristjánsson 2005; Wark et al. 2011; Roberts Kingman et al. 2021). Besides the extensive study of the three-spined stickleback genome (Hänfling et al. 2026; Nickel et al. 2026), representative chromosome-level genomes are now available for most major lineages within Gasterosteidae enabling comparative analyses (Nath et al. 2021; Liu et al. 2022; Wang et al. 2024; Chen et al. 2026). However, one crucial genome that has so far been missing is that of the sea stickleback or fifteen-spined stickleback (*Spinachia spinachia*).

The sea stickleback has long been considered to occupy a phylogenetically important position for understanding the evolution of phenotypic and genomic traits of Gasterosteidae, due to its exclusively marine habitat. The family consists of the five genera *Culaea*, *Spinachia*, *Apeltes*, *Pungitius* and *Gasterosteus*. The first three genera are monotypic, while twelve species are currently recognized within *Pungitius* and six within *Gasterosteus* (Fricke et al. 2026). Morphology, habitat, behavior and mitochondrial gene-based analyses placed *Spinachia* as the sister lineage to all other members of Gasterosteidae (McLennan 1993; McLennan and Mattern 2001; Mattern 2004). In contrast, according to molecular phylogenies inferred from 11 nuclear genes (Kawahara et al. 2009) and 1,734 single-copy orthologous genes (Liu et al. 2022), *Culaea* and *Pungitius* form a clade which is the sister to the clade of *Spinachia* and *Apeltes*. *Gasterosteus* is sister to all remaining stickleback genera.

The sea stickleback has a range of unique morphological and ecological traits compared to other members of Gasterosteidae. It is more elongated (Figure 1a) and grows to larger size than other sticklebacks (around 20 cm, compared with a maximum of around 10 cm in other stickleback genera) (Wootton 2023), possibly reflecting habitat specialization or developmental differences (Aguirre et al. 2016; Wootton 2023). It occurs in eelgrass beds and macroalgae-covered reefs in marine and brackish habitats mostly in the Northeast Atlantic (Gross 1978; Wootton 2023). It is the only stickleback species that does not frequent freshwater (Wootton 2023), although it enters brackish water in the Baltic Sea.

**Figure 1:**
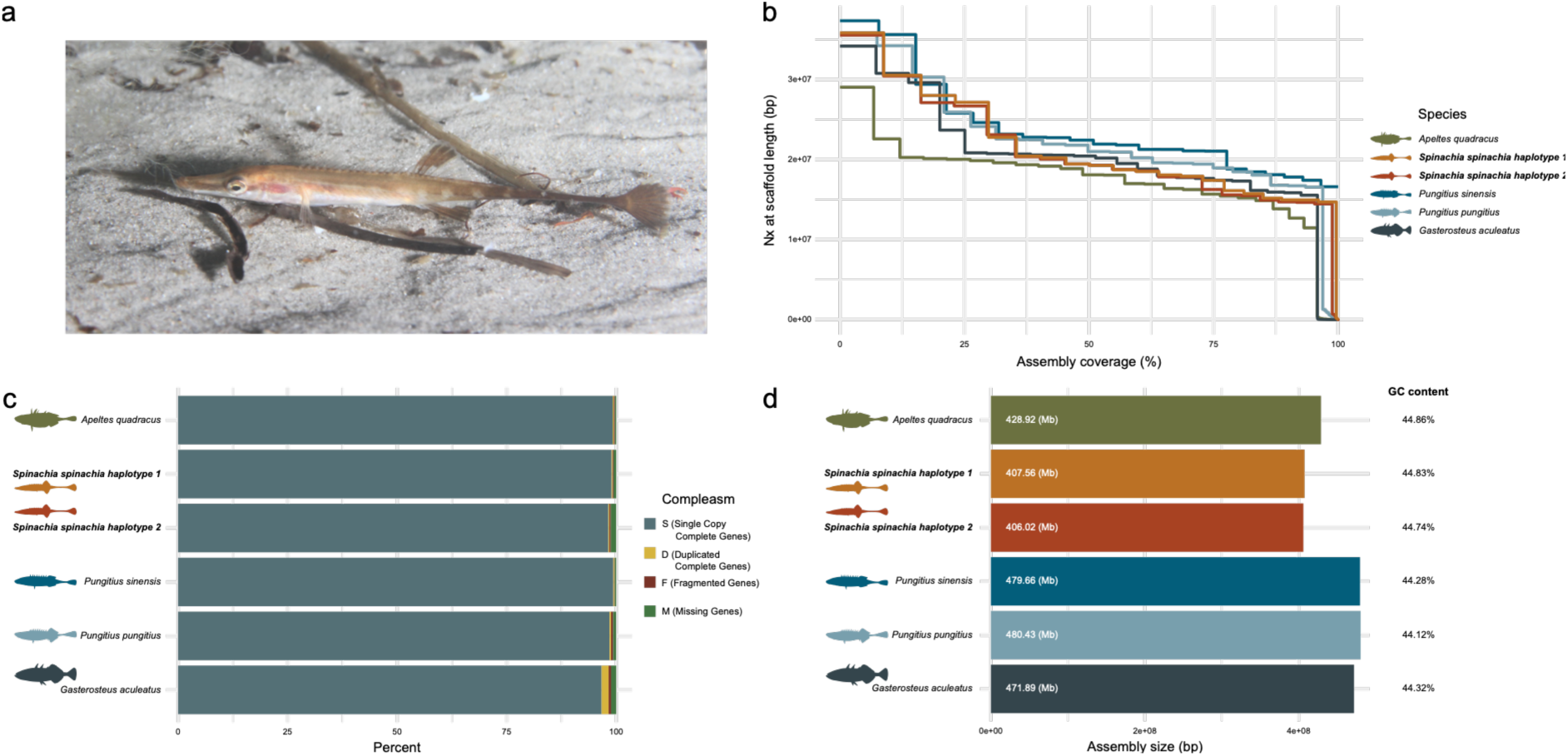
Genome assembly quality of sea stickleback (*Spinachia spinachia*) haplotypes in comparison to other Gasterosteidae assemblies. (a) A sea stickleback in its environment in Helsingør, Denmark. (b) Contiguity of the two *S. spinachia* haplotypes compared to all other published chromosome-level genomes for Gasterosteidae. The stair plot shows the percent of the genome assembly that is covered (x-axis) against corresponding Nx scaffold length (y-axis), which is the length of the shortest scaffold required to the indicated coverage percentage of the total assembly length. (c) Completeness of the *S. spinachia* genome assembly compared to other sticklebacks measured with compleasm using conserved orthologous genes defined in the actinopterygii_odb12 database. (d) Comparison across Gasterosteidae for assembly size and GC content.

Previous studies have shown that chromosome numbers vary within Gasterosteidae (2n=40–46), yet the ancestral chromosome number remains unresolved (Chen and Reisman 1970; Urton et al. 2011; Q. Li et al. 2022; Liu et al. 2022; Chen et al. 2026). The lowest number of chromosomes within Gasterosteidae is found in *S. spinachia* with 40 chromosomes, whereas its sister species fourspine stickleback (*Apeltes quadracus*) has 46 chromosomes (Liu et al. 2022). *Pungitius* has 42, while its sister species *Culaea* has 46 chromosomes, while the sister lineage to all others, *Gasterosteus*, has 42 chromosomes (Liu et al. 2022). The sister group to Gasterosteidae, represented by the tube-snout (*Aulorhynchus flavidus*, family Aulorhynchidae) has 46 chromosomes (Q. Li et al. 2022). A previously proposed evolutionary scenario includes a higher number of chromosomes (2n=46, (Chen et al. 2026)) in the ancestor of the family, which then got reduced three times, once in *Gasterosteus*, once in *Pungitius*, and once in *Spinachia* (Liu et al. 2022; Chen et al. 2026).

Studies of chromosomal rearrangements in Gasterosteidae have identified two chromosomal fusion events in *G. aculeatus* and two chromosomal fusions in *Pungitius* based on the ninespine stickleback (*P. pungitius*) and the Amur stickleback (*P. sinensis*) (Liu et al. 2022; Chen et al. 2026). Interestingly, one of these fusions was found to involve the same two homologous chromosomes of the other studied species, *A. quadracus* and *Aul. flavidus*, hence representing two independent events of the same fusion in *Gasterosteus* and *Pungitius* (Liu et al. 2022; Chen et al. 2026). Reconstruction of ancestral chromosomes also concluded that the two homologous chromosomes are fused convergently in the two genera *Pungitius* and *Gasterosteus* instead of being an ancestral fusion (Chen et al. 2026).

However, ancestral state reconstruction can change with additional taxa. The reduction in chromosome number in *S. spinachia* represents a third independent event of a change in chromosome number and may therefore enlighten the chromosome evolution in sticklebacks.

Here, we present a diploid chromosome-level genome assembly of *S. spinachia*. We use this new genomic resource to characterize chromosome fusion and fission events across Gasterosteidae and to test whether common chromosome fusions reflect convergent evolution or inheritance from ancestral karyotypes.

## Methods

### Sample acquisition, extraction, and sequencing

Two adult sea sticklebacks (specimen codes Spinspin1 and Spinspin2) were collected by push net on 2021-08-27 during a marine biology class in Kronborg Bay, Helsingør, Denmark and one individual (Spinspin4) was collected by beach seine on 2022-03-31 by the Øresund Aquarium in Kronborg Bay, Helsingør, Denmark. A physical voucher (from individual Spinspin4) was deposited at the Natural History Museum of Denmark’s (catalog number NHMD1615890, ZMUC P832248). Animals were killed using MS-222 after which tissue samples were flash frozen in liquid nitrogen. The specimens were collected under a permit (21-450C from) to the Natural History Museum Denmark.

The specimen Spinspin1 was chosen as a reference for the genome assembly. PacBio HiFi sequencing was done at the Yggdrasil Eukaryotic Reference Genome Facility at the University of Copenhagen as described in (Kliver et al. 2025). High molecular weight (HMW) DNA was extracted from 17 mg of muscles and internal organs from individual Spinspin4 using MagAttract HMW DNA Kit (Qiagen) and following the manufacturer’s protocol. One PacBio 8M wells SMRT cell was loaded and sequenced on a Pacific Biosciences SEQUEL IIe platform.

Chromatin-capture libraries were built at the Metazoa Phylogenomics Lab, Institute of Evolutionary Biology, Barcelona, Spain using the Arima HiC kit (Arima Genomics, San Diego, CA, USA) from individual Spinspin1. Sequencing of HiC libraries was done on an Illumina NovaSeq 6000 system at the Department of Biology, University of Florence, Sesto Fiorentino (FI), Italy.

To assist genome annotation, RNA library preparation was performed for four tissues (gills, brain and eyes, internal organs, and pelvic fins and muscle) collected from individual Spinspin2 at the University of Antwerp, Belgium. RNA was extracted using the Quick-RNA Miniprep Plus Kit (Zymo Research), following the manufacturer’s instructions. RNA integrity and concentration were assessed using RNA ScreenTapes with a TapeStation 4150 (Agilent Technologies) and a Qubit Fluorometer (Thermo Fisher Scientific, Waltham, MA, USA), using the Invitrogen Qubit RNA High Sensitivity assay kit. RNA samples had RINe values ranging from 7.7 to 9.5 (pelvic fins and muscle 9.5, internal organs 7.7, brain and eyes 8.9, and gills 9.3). RNA library preparation was carried out using a polyA capture protocol with the Illumina Stranded mRNA Prep kit, following the manufacturer’s instructions and using MagBio HighPrep PCR magnetic beads for library cleanup. Library fragment size distribution and concentration were assessed using High Sensitivity D5000 ScreenTapes with a TapeStation 4150 (Agilent Technologies) and a Qubit Fluorometer (Thermo Fisher Scientific, Waltham, MA, USA), respectively. RNA-seq libraries were sequenced on an Illumina NovaSeq 6000 system at the Department of Biology, University of Florence, Sesto Fiorentino (FI), Italy.

### Genome assembly

The diploid genome was assembled using the AssemblyBrute v0.1 pipeline (Kliver 2023) following the VGP assembly approach version 2 (Larivière et al. 2024) with additional quality control steps, as described in (Kliver et al. 2025). Briefly, the assembly process included the following procedures: quality control and read filtering using FastQC v0.11 (Andrews 2010), Cutadapt v3.4 (Martin 2011) and NanoPlot v1.41.6 (De Coster and Rademakers 2023), screening for contamination in reads using Kraken2 v2.1.3 (Wood et al. 2019), genome size estimation using Meryl v1.4 (Rhie et al. 2020) and GenomeScope2 v2.0 (Ranallo-Benavidez et al. 2020), phased contig assembly using Hifiasm v0.19.5-r587 (Cheng et al. 2022), identification and removal of contamination in contigs using FCS-GX v0.4.0 (Astashyn et al. 2024), purging of haplotypic duplications using purge_dups v1.2.5 (Guan et al. 2020), Hi-C scaffolding using YaHS v1.2a1 (Zhou et al. 2023), independent manual curation of haplotypes using JuiceBox (Dudchenko et al. 2018), and gap closing using SAMBA v4.1.0 (Zimin and Salzberg 2022).

### Genome annotation

To annotate repetitive regions in the genome, RepeatModeler v2.0.5 (Flynn et al. 2020) was run on haplotype 1 to identify repeat families present in the genome. Due to the resulting large number of unknown consensus sequences, TEclass2 (Bickmann et al. 2025) was used to classify these into one of the 16 families of the pre-trained model. Only classifications with a confidence of at least 80% were assigned to the RepeatModeler library. These *de novo* consensus sequences were combined with known consensus sequences from Dfam (Storer et al. 2021) for *Danio rerio* and from fishTEDB (Shao et al. 2018) for all available bony-fish species. The combined library was used to run RepeatMasker v4.1.5 (Smit et al. 2015) with default parameters for both haplotypes.

Gene annotation was performed in two different ways. EGAPx (NCBI 2024) was used to generate a *de novo* annotation, whereas TOGA2 (Malovichko et al. 2026) was used to annotate protein coding genes and infer gene orthology across all species. Annotation with TOGA2 relies on transferring protein-coding gene annotations from a reference genome, here *G. aculeatus*, to a query genome based on whole genome alignments and is described below (section Orthology inference). The main *de novo* gene annotation was generated as part of the VGP phase 1 using EGAPx, the external version of the NCBI RefSeq annotation pipeline (Formenti et al. 2026), which incorporates RNA sequencing data and related protein sequences to corroborate gene models.

### Comparison to other Gasterosteidae

In order to compare the new *S. spinachia* genome to existing stickleback genomes, we included four chromosome-level genome assemblies for Gasterosteidae, namely three-spined stickleback (*Gasterosteus aculeatus*, GCA_016920845.1 (Nath et al. 2021)), fourspine stickleback (*Apeltes quadracus*, GCA_021346845.1 (Liu et al. 2022)), ninespine stickleback (*Pungitius pungitius*, GCA_949316345.1 (Wang et al. 2024), Amur stickleback (*Pungitius sinensis*, GCA_049949165.1 (Chen et al. 2026)), and included four outgroups within Cottoidei, namely tube-snout (*Aulorhynchus flavidus*, Aulorhynchidae, available at https://datadryad.org/stash/dataset/doi:10.5061/dryad.1c59zw3w3, (Q. Li et al. 2022)), sablefish (*Anoplopoma fimbria*, Anoplopomatidae, GCA_027596085.2 (Flores et al. 2023)), limp eelpout (*Melanostigma gelatinosum*, Zoarcidae, GCA_949748355.1 (Bista et al. 2024)) and long-spined bullhead (*Taurulus bubalis*, Cottidae, GCA_910589615.1 (Potter et al. 2021)) (Table S1).

### Assembly and annotation statistics

To assess the contiguity of the new genome in comparison to the other included genome assemblies, the assemblathon_stats.pl script (Bradnam et al. 2013) was used to calculate statistics on N50, genome size, GC content and other metrics for both contigs and scaffolds. Nx was calculated using all ordered chromosome sizes. Compleasm v0.2.7 (Huang and Li 2023) was run in genome mode to assess assembly completeness based on the presence and completeness of highly conserved genes, using actinopterygii_db12 (Tegenfeldt et al. 2025) as the lineage. Additionally, compleasm was run in protein mode on the identified transcripts for both annotation methods to assess annotation completeness.

### Orthology inference

In order to infer orthologs across all available sticklebacks and selected outgroup species, TOGA2 (Malovichko et al. 2026) was run with *G. aculeatus* as the reference genome.

Therefore, all genomes were repeat masked using RepeatModeler v2.0.5 (Flynn et al. 2020) and RepeatMasker v4.1.6 (Smit et al. 2015) and additional low complexity and tandem repeats were identified and masked using WindowMasker v1.0 (Morgulis et al. 2006) and tandem repeats finder v4.09 (Benson 1999) with a maximum length of 3 Mbp. The masked genomes were then each aligned to the reference with LASTZ v1.04.41 (Harris 2007) using the parameters K=2400, L=3000, Y=9400, H=2000, and the LASTZ default scoring matrix. Local alignments were converted to chain format with axtChain (Kent et al. 2003) using default parameters except for linearGap=loose. RepeatFiller (Osipova et al. 2019) with default parameters was used to improve repetitive parts of the alignment and chainCleaner (Suarez et al. 2017) with default parameters except for minBrokenChainScore=75,000 and - doPairs was used to improve alignment specificity. After calling splice sites with spliceAI v1.3.1 (Jaganathan et al. 2019), TOGA2 was run (Malovichko et al. 2026). The output provided the basis for analyzing orthologous genes. Additionally, statistics based on genes identified by TOGA2 can be compared to genes identified by the VGP genome annotation based on transcriptomic data.

### Phylogenetic inference

To place the new *S. spinachia* genome in a phylogenetic context, we reconstructed trees using concatenation and coalescent-based frameworks based on coding regions of all Gasterosteidae and the four outgroup species. The alignments and trees per gene were created with an in-house automated pipeline (Bein et al. in prep). This automated pipeline takes a list of transcripts, a list of species and the TOGA2 reference species and extracts all one-to-one orthologs for given transcripts from all species, creates an alignment for each transcript, and builds a gene tree. One transcript was selected per gene. Priority was set in the following order: transcripts with a higher number of species having this transcript classified as fully intact, the transcript having a higher average percentage of identity and the transcript having a longer coding sequence. Alignments for each of the transcripts were performed with prank v250331 (Löytynoja 2014) and cleaned using an in-house script. Gene trees were built with iqtree3 v3.0.1 (Wong et al. 2026) with minimum branch length of 0.001 and 1,000 ultrafast bootstrap replicates. This resulted in nucleotide alignments and gene trees for 19,156 genes with at least four species present.

For the maximum likelihood inference, alignments were concatenated and partitioned per gene using AMAS (Borowiec 2016). This served as an input for iqtree3 (Wong et al. 2026), which was run with model finder (-m MFP) for 1,000 bootstrapping iterations (-B 1000). One of the chosen outgroups (*Anoplopoma fimbria*) was used to root the tree based on previous phylogenomic reconstructions (Ghezelayagh et al. 2022). For the coalescent-based approach, weighted Astral v1.25.3.8 (Zhang and Mirarab 2022) was run using the gene trees including branch lengths and bootstrap support as an input, again rooting the tree on *An. fimbria*.

### Chromosome synteny analysis

To study chromosomal rearrangements across Gasterosteidae, all genes that were identified by TOGA2 as one-to-one orthologs and classified as fully intact (FI), intact (I), partially intact (PI), or uncertain loss (UL) were extracted from the TOGA2 annotation bed file for each species. These present genes with high confidence for tracing them between species. For the TOGA reference, initially all genes were chosen. In further filtering, only one transcript per gene was kept, and genes lacking one or more species were removed in order to have each gene represented across the entire synteny plot. The positions of these 15,148 remaining one-to-one orthologous genes were then used to visualize genome synteny. This clearly showed the differences in chromosome number between species, while also allowing to visually inspect other translocations and inversions. The draw.linear function from syntenyPlotteR (Quigley et al. 2023) was used for plotting. Only chromosomes with at least 50 genes were included to reduce noise of small unplaced scaffolds. Chromosomes were sorted and flipped to maximize alignment with orthologous chromosome segments between species.

### Ancestral chromosome reconstruction

In order to better understand when the observed rearrangements likely happened in the evolutionary history of Gasterosteidae and relatives, we reconstructed the ancestral chromosomes along the phylogeny using AGORA v3.1 (Muffato et al. 2023). Like for the synteny analysis, the one-to-one orthologs were used as the input for the species.

Additionally, phylogenetic relationships were incorporated based on the tree topology obtained above. To infer the orthologs present in each internal node of the tree, orthology groups were created based on one-to-one orthologs present for all descendants of that node. AGORA was run in generic multi-pass mode, which automatically optimizes parameter settings. Resulting ancestral chromosomes were plotted using syntenyPlotteR to visually inspect rearrangements across the tree.

## Results and discussion

### Diploid chromosome-level genome assembly

Here we present the first reference genome for the sea stickleback or fifteen-spined stickleback (*Spinachia spinachia*, Figure 1a). The assembly was generated from 2.21 million PacBio HiFi reads (57.6x coverage) and 76.8 million Hi-C reads (Hi-C contact maps in Figure S1). Both haplotypes contained 20 chromosomal scaffolds each, reflecting the expected chromosome number according to the published karyotype (Liu et al. 2022). The total assembly size was 407.6 Mb and 406.0 Mb, for haplotype 1 and 2 respectively. Both haplotypes showed consistently good and comparable statistics, with high N50 values of 19.45 Mb and 19.30 Mb (Figure 1b, Table S1), and 98.96% and 98.22% completeness (actinopterrygii_odb12 database, 7,207 orthologous genes) in single copy BUSCO genes (Figure 1c, Table S1).

The sea stickleback genome assembly is slightly smaller than all other previous stickleback genomes (Figure 1d, Table S1). While *G. aculeatus* and *Pungitius* species have larger genome sizes (471-480 Mb), *S. spinachia* and its sister species *A. quadracus* have smaller genome sizes (408 and 429 Mb, respectively), possibly indicating genome size reduction in the ancestor of *Spinachia* and *Apeltes*.

Repeat annotation of the genome shows a rather small number of repetitive elements around 18.32%, most of which are transposable elements (TEs) (Table 1, Table S2). LINEs are the most prominent TE class, followed by DNA transposons, unclassified TEs and LTR elements. Other stickleback genomes similarly show few repeats with 16.99% identified in *G. aculeatus* and slightly higher numbers compared to *S. spinachia* in *P. sinensis* and *P. pungitius* with 24.41% and 32.77%, respectively (Wang et al. 2024; Chen et al. 2026; Hänfling et al. 2026). Especially, the number of identified DNA transposons is higher in the two *Pungitius* species with 10.31% and 14.8% compared to only 4.04% in *S. spinachia*.

**Table 1:** Genome assembly and annotation statistics for both haplotypes of the sea stickleback (*Spinachia spinachia*) genome. BUSCO completeness on genome and gene level was assessed using the actinopterygii_odb12 lineage using compleasm. Haplotype 2 was not annotated with the VGP pipeline, hence there are no statistics. TOGA2 annotation was performed with the three-spined stickleback (*Gasterosteus aculeatus*) as the reference. Abbreviations: S: single copy complete BUSCO genes, D: duplicated complete BUSCO genes, F: fragmented BUSCO genes, M: missing BUSCO genes, C: complete BUSCO genes as a sum of single copy (C) and duplicated (D), TEs: transposable elements, SINEs: short interspersed nuclear elements, LINEs: long interspersed nuclear elements, LTR elements: long terminal repetitive elements

| Statistic | Value haplotype 1 | Value haplotype 2 |
| --- | --- | --- |
| **Assembly statistics** |  |  |
| n contigs | 279 | 328 |
| genome size contig (Mb) | 407.54 | 405.99 |
| N50 contig (Mb) | 6.61 | 6.42 |
| n scaffolds | 38 | 52 |
| genome size scaffold (Mb) | 407.56 | 406.02 |
| N50 scaffold (Mb) | 19.45 | 19.30 |
| n chromosomes | 20 | 20 |
| compleasm S (%) | 98.96 | 98.22 |
| compleasm D (%) | 0.21 | 0.22 |
| compleasm F (%) | 0.21 | 0.32 |
| compleasm M (%) | 0.62 | 1.22 |
| GC content (%) | 44.83 | 44.74 |
| **Repeat annotation** |  |  |
| Total repeats (%) | 18.32 | 14.04 |
| TEs (%) | 16.27 | 11.67 |
| SINEs (%) | 0.86 | 1.13 |
| LINEs (%) | 5.34 | 2.38 |
| LTR elements (%) | 2.51 | 1.21 |
| DNA transposons (%) | 4.04 | 2.75 |
| Unclassified (%) | 3.53 | 4.19 |
| Simple repeats (%) | 2.07 | 2.37 |
| **Gene annotation VGP** |  |  |
| n genes VGP | 22,484 | - |
| n protein-coding genes VGP | 20,772 | - |
| n transcripts VGP | 54,641 | - |
| gene compleasm C VGP (%) | 98.78 | - |
| **Gene annotation TOGA2** |  |  |
| n protein-coding genes TOGA2 | 20,880 | 20,799 |
| n transcripts TOGA2 | 37,351 | 37,115 |
| gene compleasm C TOGA2 (%) | 97.23 | 96.6 |

The gene annotation based on the VGP pipeline identified 22,484 genes, 20,772 of which are protein-coding genes. This is similar to the 20,880 protein-coding genes identified by TOGA2 using the *G. aculeatus* gene annotation as a reference (Table 1). For haplotype 1, both annotations show high completeness as assessed by identifying more than 97% of BUSCO genes. Comparing the two annotations, the resulting number of genes is very similar with 20,772 genes in the VGP annotation and 20,880 identified by TOGA2.

Overall, the newly generated *S. spinachia* genome and its annotations of repeats and genes are of high quality and present a valuable new resource for comparative genomic analyses within the stickleback family, for example implications of this lineage on the previously studied sex chromosome diversity in the group (Dixon et al. 2019; Jeffries et al. 2022; Liu et al. 2025; Cheng et al. 2026). Here, as a first analysis, we studied chromosomal rearrangements within Gasterosteidae.

### Chromosomal rearrangements across Gasterosteidae

To reconstruct chromosomal evolution in sticklebacks, we first inferred the phylogenetic placement of *S. spinachia*. Phylogenies inferred from 19,156 protein-coding loci using maximum likelihood and coalescent-based approaches agreed on the same topology with 100% support throughout (Figure 2a, Figure S2). Relationships within Cottoidei agreed with a previous study based on ultraconserved elements (Ghezelayagh et al. 2022). The topology for Gasterosteidae with *S. spinachia* as the sister species to *A. quadracus* confirms previous studies based on mitochondrial DNA and 11 nuclear genes (Kawahara et al. 2009) and using 1,734 single-copy, orthologous genes (Liu et al. 2022).

**Figure 2:**
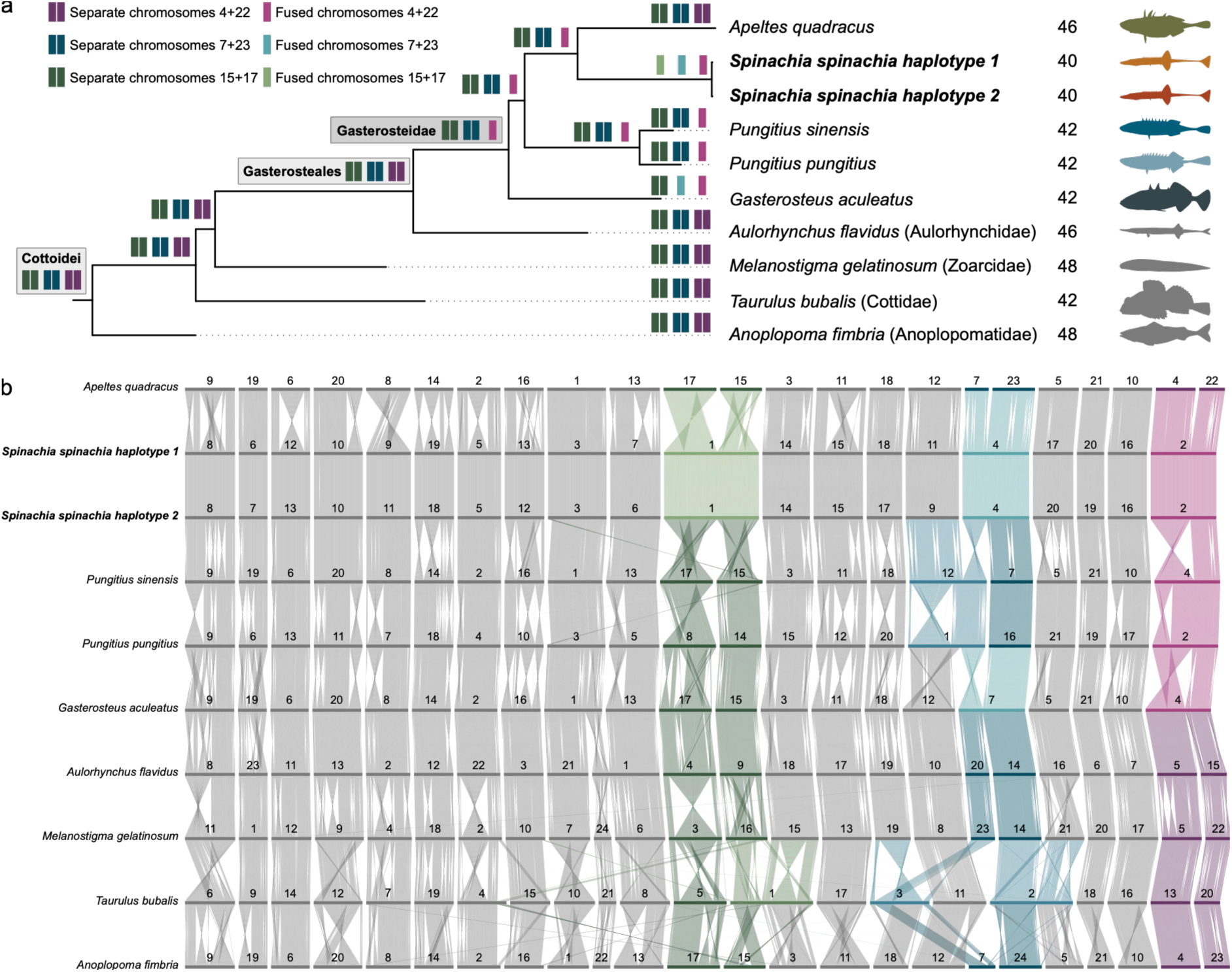
Phylogeny and chromosomal rearrangements across sticklebacks (Gasterosteidae) and outgroups within Cottoidei. (a) Three chromosomal fusions explain the low chromosome number of *Spinachia spinachia*. Phylogenetic tree based on 19,156 genes inferred using maximum-likelihood inference (coalescent-based tree was identical with posterior probabilities of 1 throughout, Figure S2). All branches had full bootstrap support. Chromosome counts are shown next to the species name.

Rectangles show chromosome reconstructions with AGORA for the chromosomes involved in the three chromosome fusions of *S. spinachia* (more detail in Figure S3). (b) Chromosomal synteny inferred from 15,148 orthologous genes, with species arranged in the order of the phylogeny.

Chromosomes that are fused in *S. spinachia* are highlighted in different colors (green: 15+17 fusion, blue: 7+23 fusion, purple: 4+22 fusion), each with the lighter shade representing a fused chromosome and the darker shade representing two separate chromosomes. If only one of the two chromosomes is involved in a fusion, this is highlighted in an intermediate color (chromosome 12 in *Pungitius sinensis*, chromosome 1 in *P. pungitius* and chromosomes 1, 2 and 3 *Taurulus bubalis*).

The karyotype of the sister species *S. spinachia* and *A. quadracus* differs by six (*Spinachia* 2n=40; *Apeltes* 2n=46). The comparison of their genomes revealed three chromosome fusion events in *S. spinachia* (Figure 2b), where chromosomes 17+15, 7+23, and 4+22 of *Apeltes* are fused into single chromosomes in *Spinachia*. The agreement with the karyotype (Liu et al. 2022) and the quality of the assemblies make it unlikely that these fusions are assembly artefacts. The chromosomes involved in these fusions appear to be often associated with translocations in other species (Figure 2b). However, in these other species, some translocations could be artifacts if the assembly was not manually curated based on Hi-C contact maps (Howe et al. 2021) and translocations should therefore be interpreted cautiously.

Fusion A, the fusion of chromosomes 17+15, was only found in *S. spinachia*. Rather than representing a simple end-to-end fusion, Fusion A appears to have been accompanied by a number of inversions and translocations (Figure 2b). These are, however, confined to the ancestral chromosomes 17 and 15 and hence preserve the ancestral chromosome gene content. One end of chromosome 17 of *Apeltes* is translocated to the opposite end in an inverted direction compared to chromosome 1 in *S. spinachia*. This could mean that instead of one of the ends, a central region on the homologous chromosome of *Apeltes* is located at the fusion point in *Spinachia* (Figure 2b). Chromosome 15 in *Apeltes* also shows several inversions and translocations compared to *Spinachia*, such that neither terminal region of *Apeltes* corresponds to the fusion point in *Spinachia*. Rearrangements within these homologous chromosomes are present in many of the other species, showing that there is flexibility in the gene order across Cottoidei.

Interestingly, the two other fusion events in *S. spinachia* (4+22 and 7+23) were found to be identical to the fusions previously described in *G. aculeatus* (Figure 2a). Fusion B, the fusion of chromosomes 4+22, was additionally found in both studied *Pungitius* species. In *S. spinachia*, chromosomes 4+22 likely fused end-to-end compared to the *Apeltes* chromosomes (Figure 2b). The same rearrangement is observed in *G. aculeatus*. In contrast, in *Pungitius* the other end of *Apeltes* chromosome 4 seems to be fused to chromosome 22. This means that both ends of chromosome 4 of *Apeltes* appear to be fused to the same end of chromosome 22 or the fusion could be followed by an inversion of one of the potential chromosome arms (Figure 2b). There seems to be an additional large inversion in *P. pungitius* within the part corresponding to *Apeltes* chromosome 4 (Figure 2b). This alternative fusion orientation in *P. pungitius* relative to *G. aculeatus* has been previously shown (Liu et al. 2022).

The phylogenetic distribution of Fusion B (4+22) allows two evolutionary hypotheses: (i) three independent fusion events in *Pungitius*, *Gasterosteus*, and *Spinachia*, or (ii) a single fusion in the ancestor of Gasterosteidae followed by a secondary fission in *Apeltes*. The first hypothesis was favored in previous studies (Q. Li et al. 2022; Chen et al. 2026), although *Spinachia* was not considered in these studies due to lack of data. Liu et al. (2022), on the other hand, acknowledged the possibility of an ancestral fusion, but concluded, in absence of additional data, that the fusions likely occurred independently.

Our results favor the second hypothesis. The presence of the same 4+22 fusion in *S. spinachia* makes an ancestral fusion followed by fission in *Apeltes* the most parsimonious explanation. Consistent with this explanation, the ancestral chromosome reconstruction using AGORA predicted that chromosomes 4 and 22 were fused in the ancestor of all Gasterosteidae (Figure 2a, Figure S3). Our reconstruction also suggests that the ancestral chromosome number of Gasterosteidae was not 2n=46 chromosomes (Chen et al. 2026) but instead lower at 2n=44 (Figure 2a, Figure S3).

Fusion C of chromosomes 7+23, was identified in both *G. aculeatus* and *S. spinachia*. In contrast to the ancestral 4+22 fusion, ancestral state reconstruction indicated that the 7+23 fusion arose independently in the two lineages (Figure 2a, Figure S3). This represents a striking example of convergent chromosomal evolution between *Spinachia* and *Gasterosteus*. The repeated fusion of the same chromosome pair in distantly related lineages suggests that certain chromosomes may be predisposed to rearrangement.

Notably, chromosome 7 is also involved in the fusion with chromosome 12 in *Pungitius*. Similarly, the distantly related long-spined bullhead (*Taurulus bubalis*) exhibits three fused chromosomes, including homologs of stickleback chromosomes 7, 15, and 23 (*A. quadracus* nomenclature; Figure 2b). Independent fusions involving the same chromosome pairs have also been reported in salmon and cod icefishes (Sutherland et al. 2016; Auvinet et al. 2020). Such recurrent fusions may reflect selection favoring particular chromosomal organizations or shared structural properties of chromosomes such as homologous or satellite repeat sequences around fusion points (Hartmann and Scherthan 2004; Yin et al. 2021; X. Li et al. 2022; Kuang et al. 2026). In butterflies, recently fused chromosomes contain large amounts of repetitive elements around fusion points, whereas these signatures become less apparent in older fusions (Wright et al. 2024). In support of the relevance of repetitive elements around fusions, we found that chromosomes 1 to 4 of *S. spinachia* showed elevated densities of repetitive elements near their central regions (Figure S4). Chromosomes 1, 2 and 3 are products of fusion events, suggesting that repetitive sequences may contribute to fusion formation or stabilization.

In summary, we found that the low chromosome count in *S. spinachia* is caused by three chromosomal fusions. Interestingly, we argue that these fusions show different patterns: Fusion A (17+15) is lineage-specific to *S. spinachia*, Fusion B (4+22) is now supported as being an ancestral fusion in Gasterosteidae, while Fusion C (7+23) is a convergent fusion in *S. spinachia* and *G. aculeatus*.

One remaining data gap is the absence of a chromosome-level genome for the Brook stickleback (*Culaea inconstans*), which possesses 46 chromosomes (Chen and Reisman 1970) and is the sister species to *Pungitius* with 42 chromosomes (Liu et al. 2022). *Culaea* hence may have split two ancestral chromosomes. Similar to the case of *Apeltes* and *Spinachia* investigated here, a genome assembly for *Culaea* could thus be insightful to study the chromosome reduction in *Pungitius*. Identifying the homologous chromosomes or potential additional rearrangements will further help to distinguish between alternative scenarios for the origin and reversal of several chromosome rearrangements identified here.

## Conclusion

The new diploid sea stickleback genome fills a crucial gap for studying Gasterosteidae, a family that encompasses the model organism *G. aculeatus*. The *S. spinachia* genome helped reinterpret the karyotypic variation present in the group. The results revealed three distinct modes of chromosomal evolution in sticklebacks. Fusion A (17+15) is unique to *Spinachia*, Fusion B (4+22) is best explained by fusion in the ancestor of Gasterosteidae with subsequent fission in *Apeltes*, and Fusion C (7+23) represents a case of convergent chromosome evolution in *Spinachia* and *Gasterosteus*. Together these results demonstrate that chromosome evolution of Gasterosteidae includes secondary fission of ancestrally fused chromosomes and convergent fusion of the same chromosome pairs. In the future, these repeated events could be further investigated to understand whether the changed recombination and gene linkage of fused or split chromosomes ultimately have an impact on speciation and adaptation (Guerrero and Kirkpatrick 2014; Liu et al. 2022; Yoshida et al. 2023; Diblasi and Saitou 2026; Hoff et al. 2026). These recurrent events provide a valuable system for investigating the structural properties of chromosomes facilitating or hindering fusions and fissions and their evolutionary consequences. The insights from the *S. spinachia* genome also highlight the importance of sampling coverage, and future inclusion of *Culaea inconstans* will help further resolve the history of karyotype evolution in sticklebacks.

## Data availability

The *Spinachia spinachia* genome with both haplotypes is available on NCBI with accession numbers GCA_048126635.1 for haplotype 1 and GCA_048127205.1 for haplotype 2. The associated BioProjects PRJNA1146010, PRJNA1146009 include the whole genome nucleotide and protein sequences, while PRJNA1393321 and PRJNA1393320 include the genome and transcriptome sequencing data, respectively. The sample used for genome sequencing can be found under BioSample SAMN36735485, the samples for transcriptome sequencing for different tissue types under SAMN43111165, SAMN43111166, SAMN43111167 and SAMN43111168. The VGP annotation for haplotype 1 is linked to the assembly under accession GCA_048126635.1-GB_2025_08_04 (https://ftp.ncbi.nlm.nih.gov/genomes/all/GCA/048/126/635/GCA_048126635.1_SpiSpi1_v1. hap1/) and can also be found on GenomeArk (https://genomeark.s3.amazonaws.com/index.html?prefix=downstream_analyses/EGAPx_a nnotations/fSpiSpi1_output/).

Assembly and annotation pipelines are available on GitHub (https://github.com/mahajrod/AssemblyBrute). All used scripts and data for the phylogenetic and synteny analysis will be available on GitHub upon publication.

## Acknowledgements

This work was supported by a research grant (42153) from VILLUM FONDEN to JS, and a grant from German Research Foundation to LH (HI2214/1-1, Project number: 530763738). Computation was facilitated by allocations from the Danish e-infrastructure Consortium (DeiC-KU-N2-2024080, DeiC-KU-N2-2025160) to JD and JS. This genome was generated as part of the European Reference Genome Atlas (ERGA) Pilot Project (Mazzoni et al. 2023; Mc Cartney et al. 2024). We are grateful to Shyam Gopalakrishnan (University of Copenhagen) for help on the ERGA Pilot Project. Lab work and genome assembly was made possible through the Yggdrasil project funded by Carlsbergfondet Research Infrastructure Grant (CF22-0680) to MTPG. Long-read data generation was supported thanks to the Danish National Research Foundation award (DNRF143) to MTPG. The PacBio Facility at University of Copenhagen’s Biology Department is funded by Novo Nordisk Foundation Award NNF20OC0061528 (“Copenhagen DNA Analysis Center (CoDoN)”) to Prof Søren J Sørensen. RF acknowledges support from the following sources of funding: Ramón y Cajal fellowship (grant agreement no. RYC2017-22492 funded by MCIN/AEI /10.13039/501100011033 and ESF ‘Investing in your future’), the Agencia Estatal de Investigación (project PID2019-108824GA-I00 funded by MCIN/AEI/10.13039/501100011033), the European Research Council (this project has received funding from the European Research Council (ERC) under the European Union’s Horizon 2020 research and innovation programme (grant agreement no. 948281)), the Human Frontier Science Program (grant no. RGY0056/2022) and the Secretaria d’Universitats i Recerca del Departament d’Economia i Coneixement de la Generalitat de Catalunya (AGAUR 2021-SGR00420). Illumina sequencing (RNA and Hi-C libraries) was supported by the sequencing facility of the Department of Biology, University of Florence through the Departments of Excellence programme funded by the Italian Ministry for University and Research. We thank the Antwerp University Hospital Center of Medical Genetics and Jarl Bastianen for access to sequencing library quality control equipment. We acknowledge access to the storage resources at Barcelona Supercomputing Center, which are partially funded from the European Union H2020-INFRAEOSC-2018-2020 programme through the DICE project (Grant Agreement no. 101017207). We would like to thank Alisha Ahamed, Josephine Burgin, Joana Paupério, Jeena Rajan and Guy Cochrane from the European Nucleotide Archive (ENA) for their support regarding data coordination and submission. We would like to thank Felix Shaw, Aaliyah Providence, Debby Ku, Robert P. Davey, Seanna McTaggart, and Alice Minotto for their support regarding metadata coordination and submission.

