## Supplementary tables and figures for "Sea stickleback genome reveals repeated chromosomal rearrangements in sticklebacks"

**Table S1**: Sources and statistics for all Cottoidei genomes used. Provided as a separate supplementary file in xlsx format (TableS1.xlsx)

**Table S2**: Repeat annotation of *Spinachia spinachia* haplotype 1. The table shows the summary of repetitive elements identified by RepeatMasker. The number, total length and percentage of the genome are given for each category of elements.

| | Number of elements | Length occupied (bp) | Percentage of sequence (%) |
|---|---|---|---|
| **Total bases masked** | | **74,664,490** | **18.32** |
| Retroelements | 254,350 | 35,482,369 | 8.71 |
| SINEs | 12,974 | 3,497,939 | 0.86 |
| Penelope | 0 | 0 | 0.00 |
| LINEs | 182,296 | 21,766,515 | 5.34 |
| CRE/SLACS | 0 | 0 | 0.00 |
| L2/CR1/Rex | 69,559 | 9,628,662 | 2.36 |
| R1/LOA/Jockey | 13,572 | 393,911 | 0.10 |
| R2/R4/NeSL | 14,410 | 1,109,418 | 0.27 |
| RTE/Bov-B | 28,141 | 2,761,488 | 0.68 |
| L1/CIN4 | 19,839 | 1,559,049 | 0.38 |
| LTR elements | 59,080 | 10,217,915 | 2.51 |
| BEL/Pao | 1,408 | 294,042 | 0.07 |
| Ty1/Copia | 857 | 119,277 | 0.03 |
| Gypsy/DIRS1 | 29,973 | 5,670,893 | 1.39 |
| Retroviral | 5,964 | 1,135,813 | 0.28 |
| DNA transposons | 131,689 | 16,446,293 | 4.04 |
| hobo-Activator | 41,300 | 5,623,917 | 1.38 |
| Tc1-IS630-Pogo | 19,348 | 3,835,780 | 0.94 |
| En-Spm | 0 | 0 | 0.00 |
| MULE-MuDR | 363 | 41,673 | 0.01 |
| PiggyBac | 790 | 183,441 | 0.05 |
| Tourist/Harbinger | 2,740 | 399,247 | 0.10 |
| Other (Mirage, P-element, Transib) | 15,402 | 1,293,750 | 0.32 |
| Rolling-circles | 85 | 10,790 | 0.00 |
| Unclassified | 53,088 | 14,367,635 | 3.53 |
| **Total interspersed repeats** | | **66,296,297** | **16.27** |
| Small RNA | 2,298 | 175,240 | 0.04 |
| Satellites | 691 | 72,589 | 0.02 |
| Simple repeats | 172,087 | 7,386,816 | 1.81 |
| Low complexity | 16,731 | 835,151 | 0.20 |

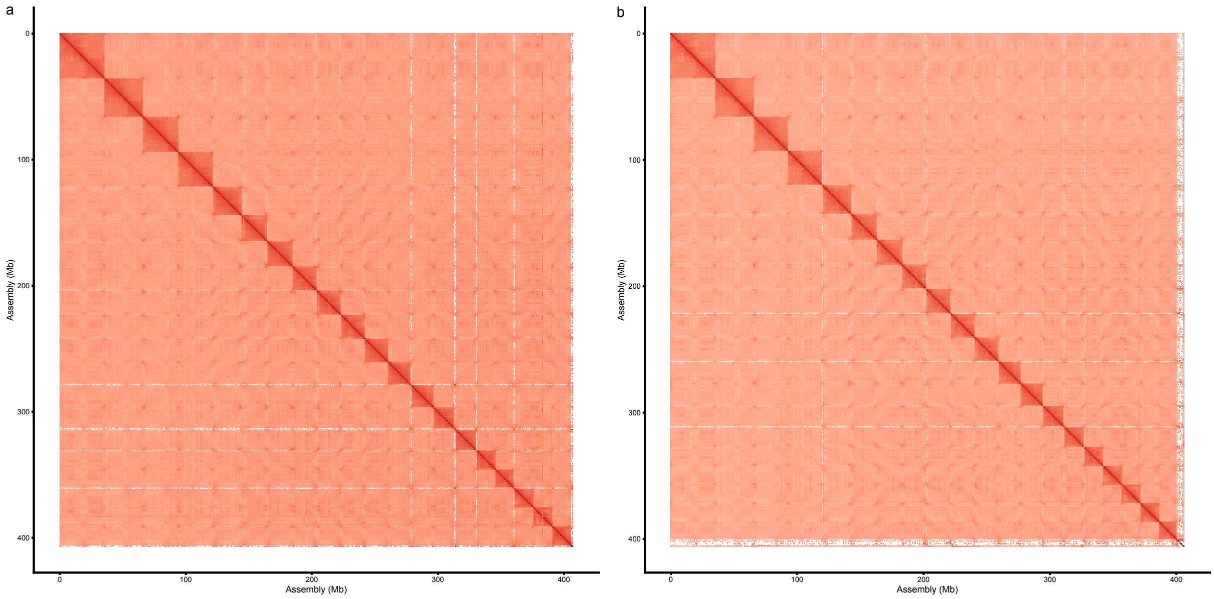


**Figure S1**: Hi-C contact map for (a) haplotype 1 and (b) haplotype 2.

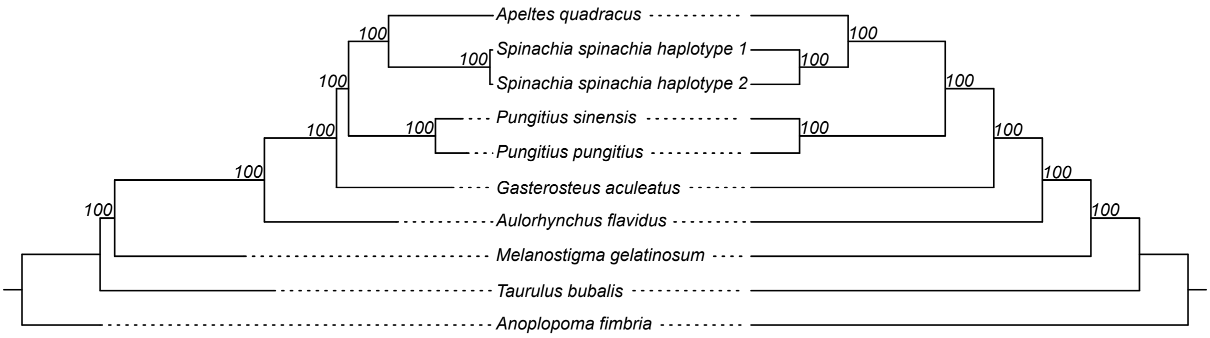


**Figure S2**: Comparison of maximum-likelihood and coalescent based trees. The tree on the left was built based on 19,156 gene alignments with iqtree3. The tree on the right was built based on 19,156 gene trees using weighted Astral. Both tree topologies are identical, and both trees show very high support values for all internal nodes, measured as bootstrap support on the left and local posterior probability in percent on the right.

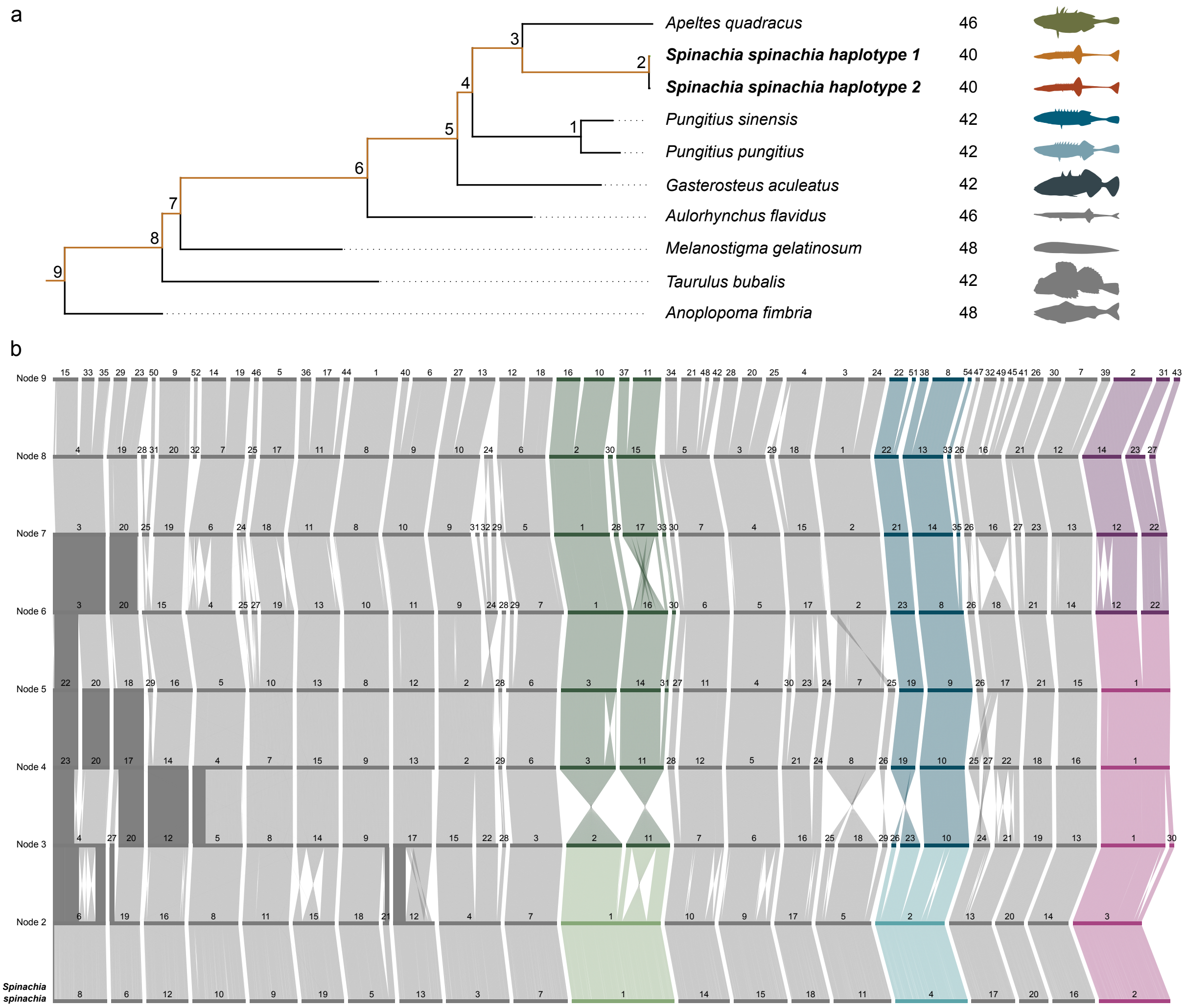


**Figure S3**: Phylogeny and ancestral chromosome reconstruction. (a) Phylogenetic tree of Gasterosteidae and four outgroup species based on 19,156 genes. The path highlighted in orange, from *Spinachia spinachia* to the root, corresponds to the nodes shown in the synteny plot in (b). Species chromosome counts are stated after the species name. (b) Synteny of ancestral chromosomes reconstructed with AGORA. Node labels correspond to panel (a). The three chromosomes that are fused in *S. spinachia* are highlighted in different colors, with the lighter color representing a fused chromosome and the darker color representing two separate chromosomes.

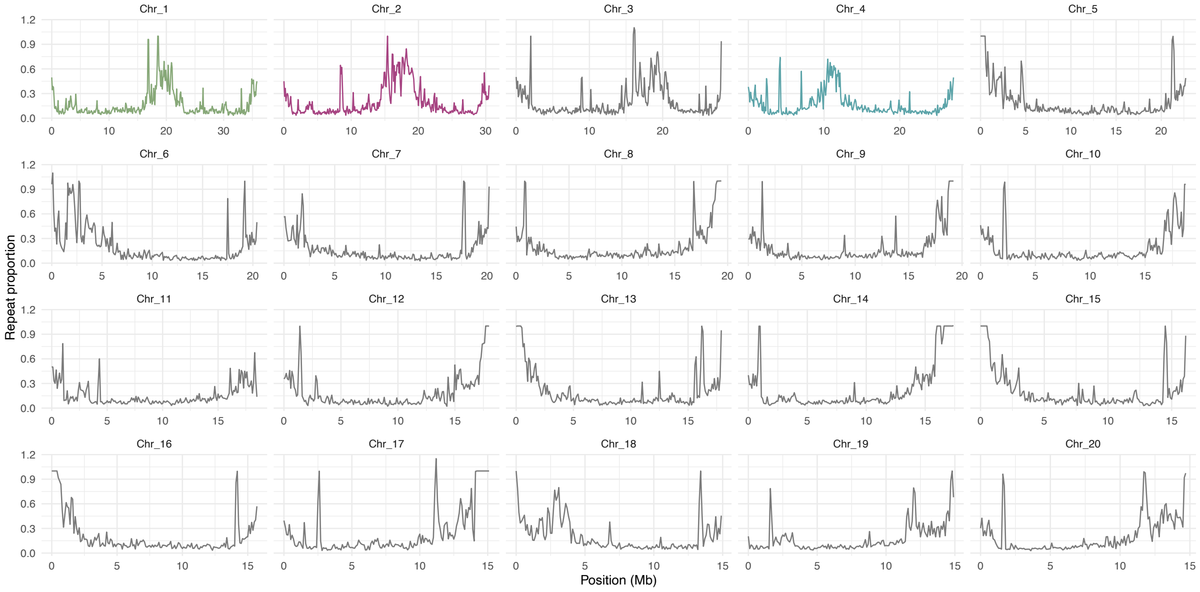


**Figure S4**: Repeat coverage per 100 kB windows for each chromosome of *Spinachia spinachia* haplotype 1. Fused chromosomes are highlighted in colors matching Figure 2 and Figure S3.
